# A Statistical Physics Approach for Disease Module Detection

**DOI:** 10.1101/2022.01.26.477756

**Authors:** Xu-Wen Wang, Dandi Qiao, Michael Cho, Dawn L. DeMeo, Edwin K. Silverman, Yang-Yu Liu

## Abstract

Extensive evidence indicates that the pathobiological processes of a complex disease are associated with perturbation within specific disease neighborhoods of the human protein-protein interaction (PPI) network (a.k.a. the interactome), often referred to as the disease module. Many computational methods have been developed to integrate the interactome and omics profiles to extract context-dependent disease modules. Yet, existing methods all have fundamental limitations in terms of rigor and/or efficiency. Here, we developed a statistical physics approach based on the random-field Ising model (RFIM) for disease module detection, which is both mathematically rigorous and computationally efficient. We applied our RFIM approach with genome-wide association studies (GWAS) of six complex diseases to examine its performance for disease module detection. We found that our RFIM approach outperforms existing methods in terms of computational efficiency, connectivity of disease modules, and robustness to the interactome incompleteness.

## INTRODUCTION

Complex diseases are believed to be caused, among other factors, by combinations of genetic alterations affecting cellular systems^1^. Integration of disease-related genomics, epigenomics, transcriptomics and other omics data with the human protein-protein interaction network (or interactome) could provide insight into the molecular mechanisms of complex diseases. A disease network module represents a group of cellular components (e.g., dysfunctional pathways, complexes or crosstalk subgraphs between pathways in a molecular interaction network) that together contribute to normal cellular function that, when disrupted, may produce a particular disease phenotype^1^. The concept of the disease module is based on the local hypothesis that if a gene or molecule is involved in a specific biochemical process or disease, its direct interactors might also be suspected to play some role in the same biochemical process or disease^2–4^. For example, proteins that are involved in the same disease show a high propensity to interact with each other^5,6^, and genes that are mutated in diseases with similar phenotypes have a significantly increased tendency to interact directly with each other^7,8^. Hence, we expect that each disease can be linked to a well-defined neighborhood of the cellular molecular interactome, i.e., a *disease module*, which is likely to include multiple disease-modifying genes that mediate various epigenetic, transcriptional and post-translational phenomena. Detection of the disease module for a pathophenotype of interest in turn can guide further experimental work towards uncovering the disease mechanism, predicting disease genes and informing drug development^9–13^.

There are many existing methods for disease module detection^14–16^. For example, ModuleDiscoverer^17^ used a heuristic-based approximation of the community structure based on the maximal clique enumeration formalism. DIAMOnD^18^ starts from a set of seed genes and prioritizes the other proteins of the interactome for their putative disease relevance. However, those methods all have fundamental limitations. For example, seed-gene based methods heavily rely on the initial selection of seed genes. As for heuristic-based methods that attempt to find optimal subgraphs within the molecular interaction network, they turn out to be computationally intractable in general.

To address the limitations of previous methods, here we develop a novel statistical physics approach for disease module detection based on the random-field Ising model (RFIM)^19–21^. We hypothesize that optimizing the “score” of the whole network rather than a particular subgraph will discover a biologically more meaningful disease module. Optimizing the score of the whole network (with appropriately assigned node weights and edge weights calculated from omics profiles) can be mapped to the ground state problem of RFIM and then solved exactly by the max-flow algorithm with polynomial time complexity^22^. The resulting disease module might not be a single connected component or a tree graph at all. Using this approach, all of the disease related subgraphs will emerge simultaneously.

To compare our RFIM approach with existing methods, we applied our RFIM approach and two representative methods to genome-wide association studies (GWAS) of six different complex diseases: asthma^23^, breast cancer^23^, chronic obstructive pulmonary disease^24^ (COPD), cardiovascular disease^25^ (CVD), diabetes^26^, and lung cancer ^23^. All SNPs were annotated to the nearest genes and gene-wise p-values were obtained by using a known approximation of the sampling distribution using MAGMA^27–29^. These gene-wise p-values were further integrated with the human interactome to identify the disease module. We found that our RFIM approach displays high computational efficiency and superior scalability. Moreover, the disease module identified by RFIM is much denser and includes more disease-associated pathways than other methods. Finally, we demonstrated clear evidence that RFIM is less sensitive to the incompleteness of the interactome than other methods.

## METHOD

We formalize the disease module detection problem as a statistical physics problem as follows. We represent the state of node (gene) *i* by a binary variable *σ_i_* = ±1, where +1 (or −1) means gene *i* is “active” (or “inactive”), i.e., belongs to the disease module (or not), respectively. The disease module is captured by the state of all genes, denoted as {*σ_i_*}. Given a graph *G* = (*V, E*), i.e., the PPI network with a node set *V* (with node weights {*h_i_*}) and an edge set *E* (with edge weights {*J_ij_*}), we aim to find the optimal gene state {*σ_i_*}, also called the ground state, that minimizes the following cost function:

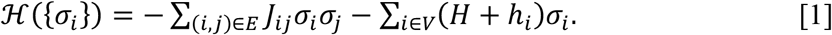

The subgraph induced by the active nodes in the ground state is the disease module.

The first term in the right-hand side of Eq. [1] represents the influence of nodes (genes) on their neighbors’ states. By defining non-negative edge weights {*J_ij_*}, we implicitly assume that neighboring nodes (genes) in the interactome tend to have the same state, i.e., they tend to belong to the disease module (or not) together. In other words, *J_ij_* > 0 favors positive correlations between the states of nodes *i* and *j*. The stronger the *J_ij_* value, the more likely nodes *i* and *j* have the same state. In the case of *J_ij_* = 0, the states of nodes *i* and *j* are totally independent. The second term in the right-hand side of Eq. [1] stands for the intrinsic preference of nodes belonging to the disease module regardless of the states of their neighboring nodes. The intrinsic preference of node *i* can be quantified as *h_i_* = Φ^−1^(1 – *p_i_*), where Φ^−1^ is the inverse normal cumulative distribution function and *p_i_* is the p-value of node (gene) *i* calculated from the omics profile (see **Fig.1a**). Note that *h_i_* is a monotonically decreasing function of *p_i_* with *h_i_* → +∞ as *p_i_* → 0 and *h_i_* → −∞ as *p_i_* → 1. Nodes with very positive *h_i_* (or equivalently, very small *p_i_*) tend to belong to the disease module.

**Figure 1:**
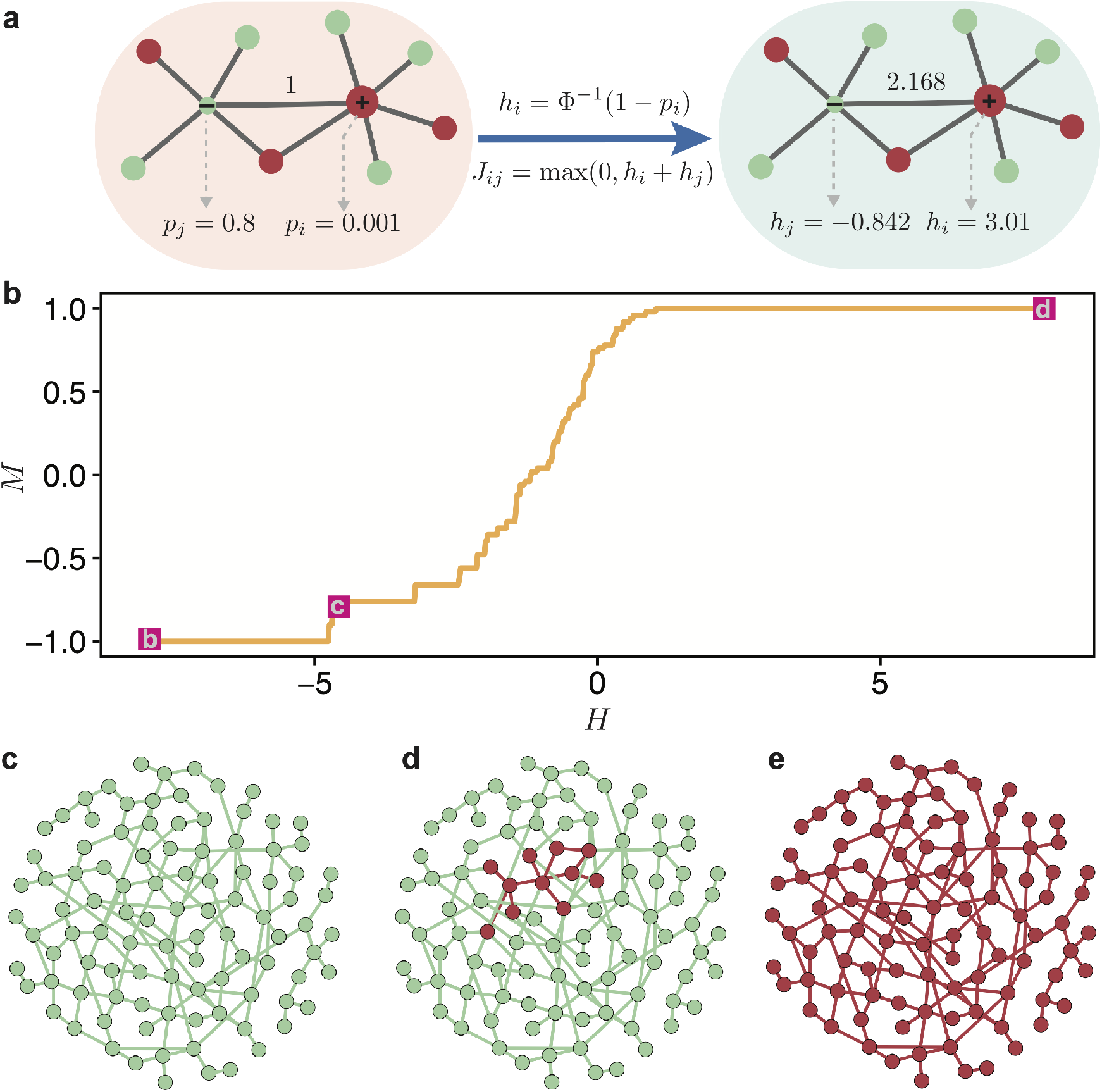
Disease module inference using statistical physics approach: a demonstration. **(a):** Schematic demonstration of the RFIM approach. A random network with *N* = 100 nodes (genes). Each gene is assigned a p-value, randomly chosen from a uniform distribution *U*(0,1). The node weights *h_i_* are then computed using the inverse normal CDF. The edge weights *J_ij_* = max (0, *h_i_* + *h_j_*) if two genes are connected in the network. Otherwise, *J_ij_* = 0. **(b):** The magnetization curve *M*(*H*) of the random network system. The magnetization 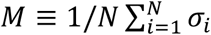 is the average state of nodes in the system. As we increase the external field *H*, more nodes (genes) will be active, i.e., *M* will monotonically increase and eventually approach 1. **(c-e):** disease module identified by RFIM at *H* = –8 (point b), *H* = –4 (point c) and *H* = 8 (point d), respectively. All the active components (highlighted in red) together can be considered as the disease module.

The parameter *H* is introduced to regulate the contributions of the two terms in Eq. [1]. Note that the value of *H* effectively controls the average state variable 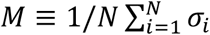 of the system, or equivalently, the disease module size given by (1 + *M*)*N*/2 (see **Fig.1b**). For example, if *H* is extremely low such that (*H* + *h_i_*) < 0 for all the nodes, then the ground state is simply {*σ_i_* = –1}, i.e., all nodes will be inactive (*M* = –1 and there is no disease module, which is a trivial solution, **Fig.1c**). With gradually increasing *H*, more nodes will be active (**Fig.1d**). Eventually, if *H* is high enough such that (*H* + *h_i_*) > 0 for all the nodes, then the ground state is {*σ_i_* = +1}, i.e., all nodes will be active (*M* = +1 and the whole interactome is the disease module, which is also a trivial solution, **Fig.1e**). Hence, controlling the parameter *H*, we can obtain a disease module with any desired size between 0 and *N*.

Note that Eq. [1] describes exactly the energy (or “Hamiltonian”) of the RFIM rooted in statistical physics, especially in the study of disordered magnets^30^. In its most general form, the RFIM applies to an ensemble of magnetic particles interacting with each other in the presence of an external magnetic field *H*. Each magnetic particle (also called spin) has two states *σ_i_* = ±1 (i.e., UP or DOWN). *h_i_*’s are local magnetic fields (due to impurities) acting on each spin and *h_i_* >0 (or < 0) favors UP (or DOWN) state of spin *i*, respectively. The coupling constants *J_ij_* > 0 capture the ferromagnetic interactions between spins *i* and *j* such that two spins next to each other will tend to align to save the energy of the system. The control parameter *H* is just the external magnetic field, acting uniformly on all the spins to align them in the direction of *H*. The average state variable (or order parameter) *M* is the magnetization of the spin system.

In our calculation, we define the edge weight *J_ij_* = max (0, *h_i_* + *h_j_*) if nodes *i* and *j* are *connected* in the interactome, and *J_ij_* = 0 otherwise. This definition of *J_ij_* is to ensure that (1) the two terms in the right-hand side of Eq. [1] are numerically comparable; (2) *J_ij_* is non-negative even in case the intrinsic preference of nodes *i* (or *j*) belonging to the disease module is extremely low (i.e., with *h_i_* or *h_j_* → −∞). Note that the disease module calculated from the active nodes in the ground state of the RFIM is not necessarily a single connected component. In fact, it might contain multiple connected components in the network *G*. To ensure the connectivity of the disease module is not too low, we modify the intrinsic preference of node *i* belonging to the disease module as 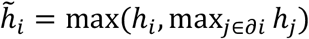, where *∂i* represents the neighborhood (i.e., the set of neighboring nodes) of node *i*.

Our RFIM approach for the disease module detection has several advantages and novelties over existing approaches: (1) Since the ground state and the *M*(*H*) curve of the RFIM can be exactly calculated with polynomial time complexity^22,31^, we can run the calculation on very large data sets, e.g., human interactome, without any difficulty. (2) Both edge weights and node weights are naturally considered. (3) The inferred disease module is not necessarily a single component or a tree graph at all.

## RESULTS

### Associations between gene significance and degree

We considered gene-wise p-values calculated from the GWAS of six complex diseases: asthma^23^, breast cancer^23^, chronic obstructive pulmonary disease^32^ (COPD), cardiovascular disease^25^ (CVD), diabetes^26^, and lung cancer^23^, respectively. For each disease, we overlaid the GWAS gene-wise p-values onto the human interactome. For the human interactome, we used two well-established ones: (1) STRING^23^; and (2) iRefIndex^34,35^. Before we identify the disease module, we examined the association between the gene-wise p-value and the gene (node) degree in the interactome. We found that those genes with low p-values (or high p-values) in GWAS typically have low degrees (or high-degree, i.e., hubs) in the interactome, respectively. Interestingly, this association is quite robust to different diseases and interactomes (see **Fig.2**). We suspected that this association might be related to the gene essentiality. For example, a previous study on the PPI of the yeast *Saccharomyces cerevisiae* found that many hub genes are essential for survival of the cell, and yeast cannot grow and multiply without them^36^. Thus, the SNPs annotated to the essential genes are less likely to be functional variants. To validate this hypothesis, we used a reference of gene essentiality constructed by genome-wide single-guide RNA library for genes required for proliferation and survival in a human cancer cell line^37^. A CRISPR score is defined as the average log2 fold-change in the abundance of all single-guide RNA targeting a given gene and of the 18,166 genes, 1,878 scored as essential. We employed this reference to dichotomize genes in GWAS to essential and non-essential. We found that the gene-wise p-values of essential genes in the GWAS of asthma, CVD, and diabetes are significantly higher than that of non-essential genes (see **Fig.S1a**). Moreover, we found that the degrees of those essential genes are significantly higher than non-essential genes over all six diseases and regardless of the interactome analyzed here (see **Fig.S1b,c**).

**Figure 2:**
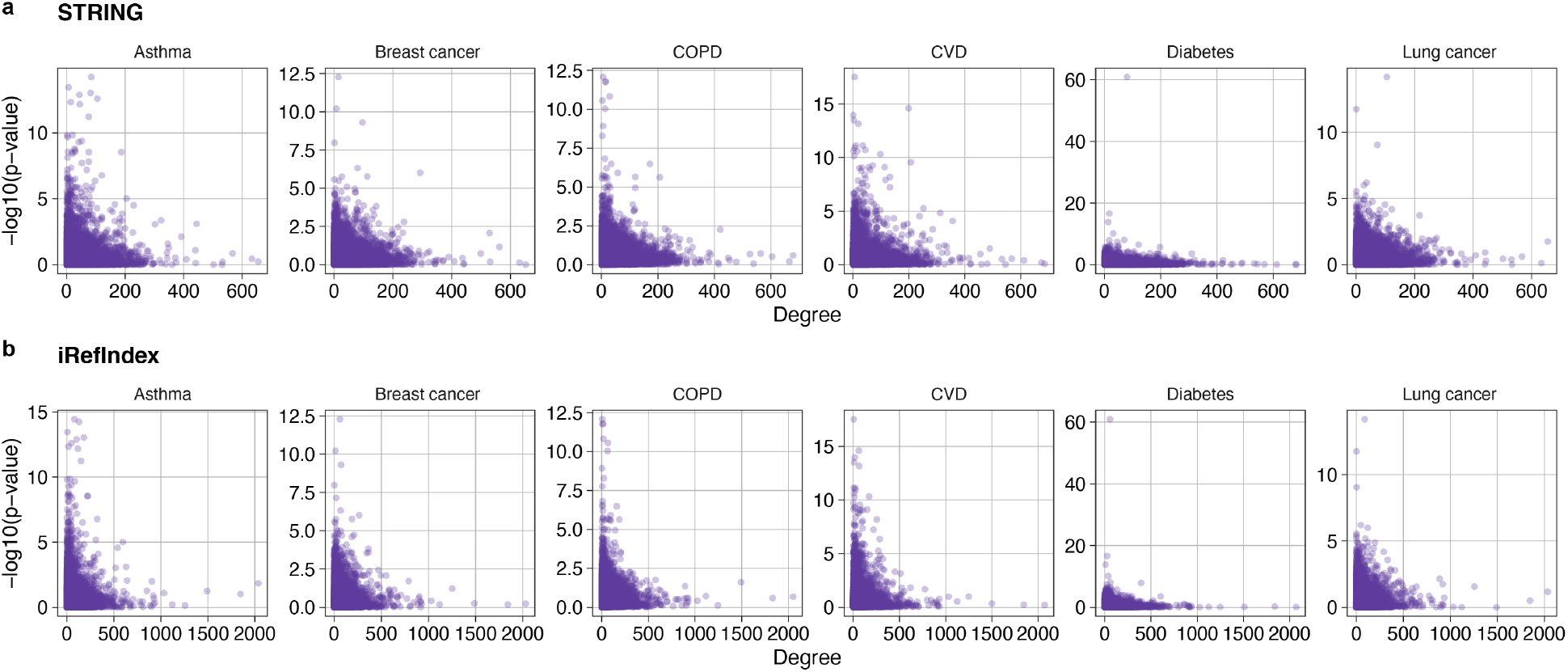
Association between gene-wise p-value and gene degree. For each phenotype, we showed the gene-wise p-value (log10 scale) vs. its degree in the interactome of **(a):** STRING and **(b):** iRefIndex.

### Computational complexity of disease module detection

After overlaying the GWAS gene-wise p-values onto the human interactome, we can identify the disease module using various methods. For our RFIM approach, we calculated the *M*(*H*) curve of the RFIM by systematically tuning *H*. The disease module is detected from the ground state of the RFIM at *H* = *H*_c_. We chose *H*_c_ to ensure the disease module reaches a desired size (i.e., consisting of 5% of the total genes in the GWAS). In DIAMOnD, the most significantly connected node (lowest p-value) is integrated into the module at each step of the iteration until the desired module size is reached^18^ (and we chose the desired module size to be 500 for each disease). Given an interactome, ModuleDiscoverer first approximates the underlying community structure by iterative enumeration of gene cliques from random seed genes. Then, the union of all significantly enriched cliques are ensembled into a large module^17^. We assessed the computational complexity of different methods measured as the average computing time across all the six diseases. We found that RFIM and DIAMOnD are the two most efficient methods, which require significantly shorter running time than ModuleDiscoverer (see **Fig.3**). Moreover, we found that RFIM shows superior scalability over other methods across different interactomes. This scalability is due to the fact that the ground state of RFIM can be exactly solved with polynomial time complexity.

**Figure 3:**
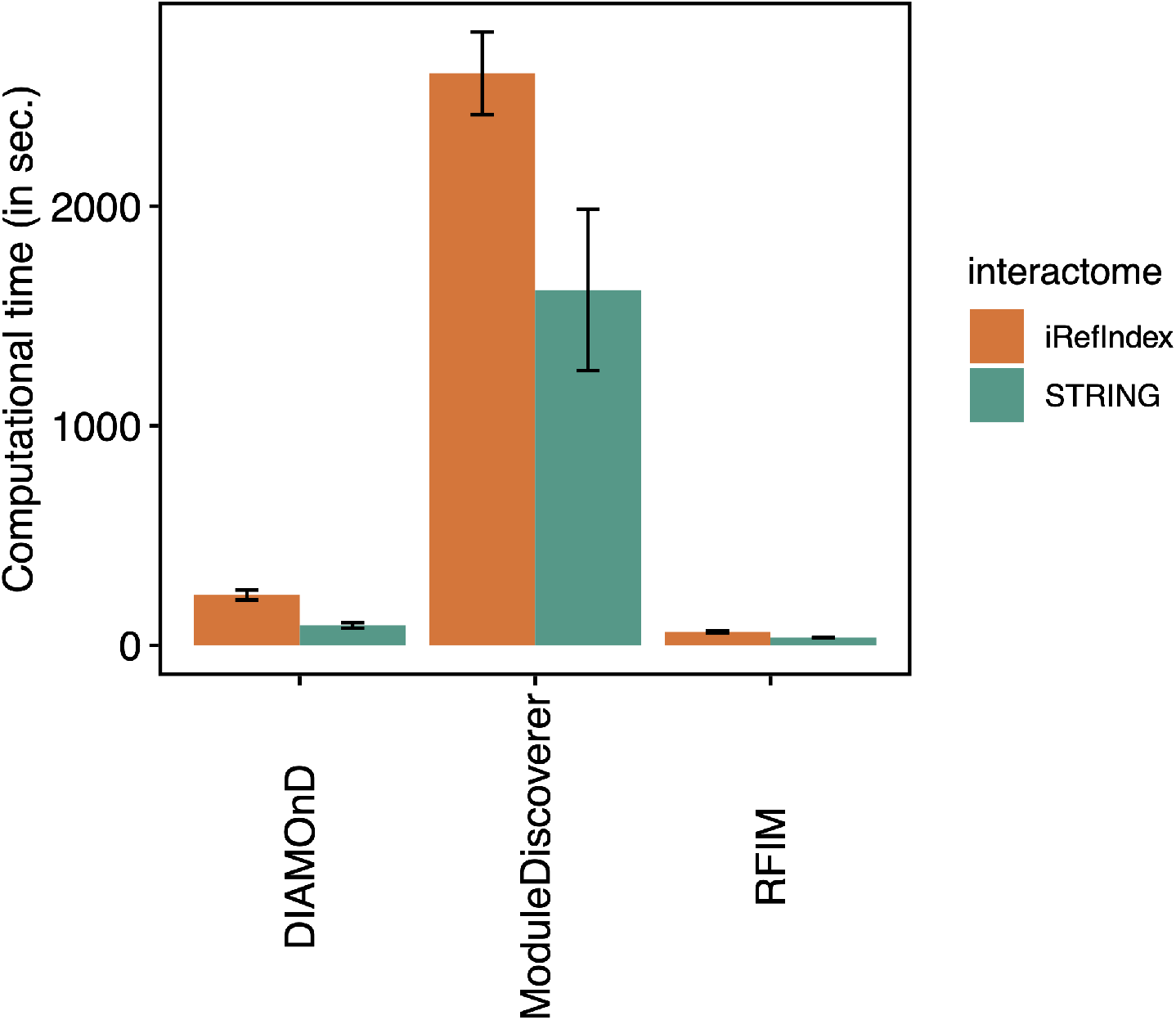
Computational complexity of each disease module detection method. Each bar represents the average computational time (in seconds) overall all six phenotypes. Error bar represents the standard derivation among 6 GWAS datasets. The computational time is measured as the running time on macOS machine with 2.4 GHz 8-Core Intel Core i9 processor.

### Connectivity of the disease module

By definition, the disease module is a specific neighborhood within the interactome. Hence, the connectivity of the disease module should not be too low. We calculated the mean degree of the disease module detected by each method, finding that the disease module identified by RFIM shows the highest mean degree in all (or two of the six) diseases using the STRING (or iRefIndex) interactome, respectively (see **Fig.4a,b**). Moreover, we found that the mean degree of disease module identified from RFIM is higher or comparative to the mean degree of the whole interactome for all the diseases analyzed in this work. Yet, the mean degree of disease modules identified by DIAMOnD (or ModuleDiscoverer) is much lower than that of the whole interactome for most diseases considered here.

**Figure 4:**
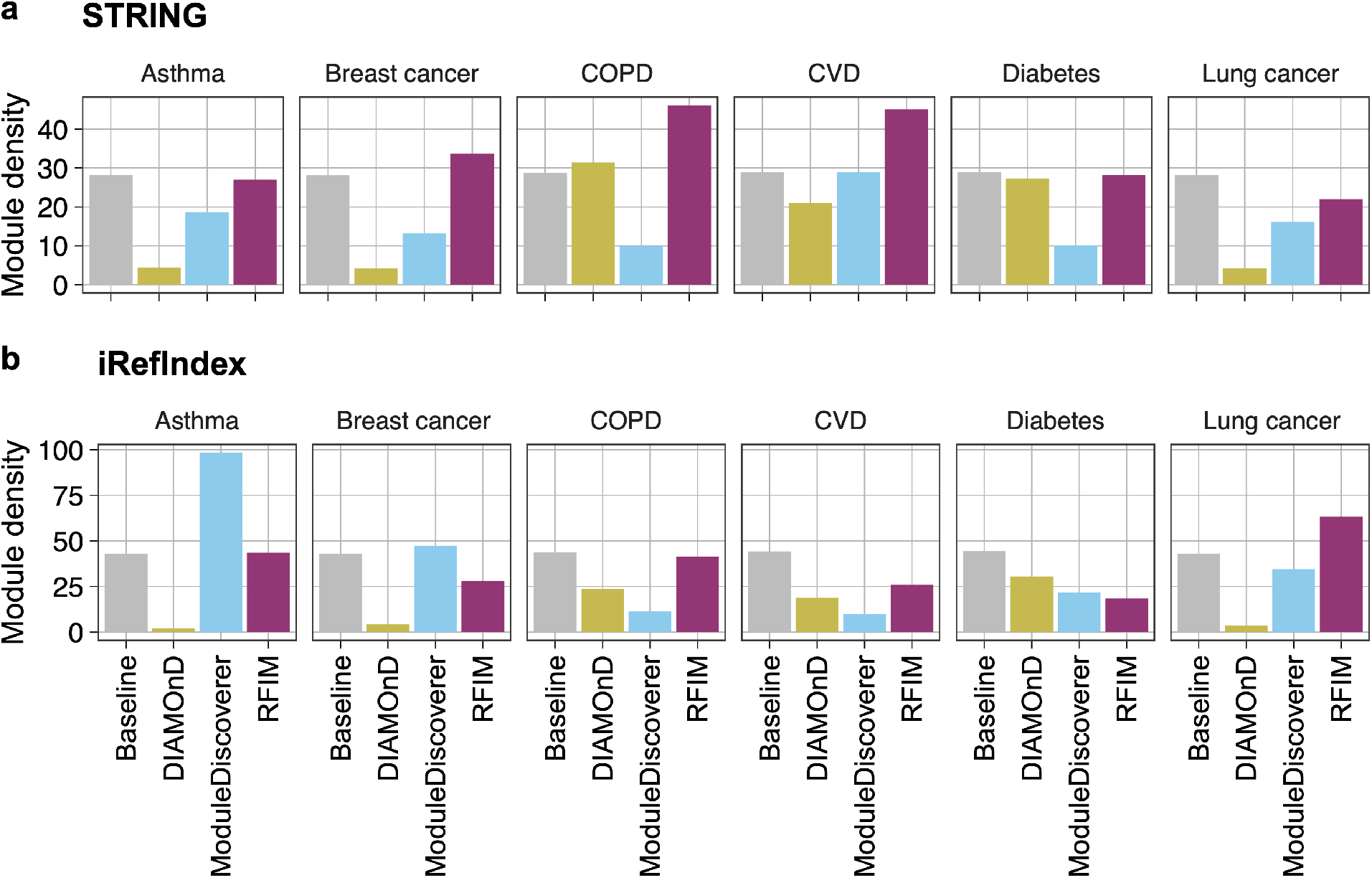
Mean degree of disease module for each phenotype and method. We applied each disease module detection method in the GWAS of each disease separately. Baseline represents the average degree of the interactome. For each disease, we chose to work on the common genes between the interactome and the GWAS, thus the average degree of the interactome in different diseases slightly varies.

### Pathway enrichment analysis

To gain mechanistic insights into the list of genes in the disease module, we performed pathway enrichment analysis. In particular, we employed the ReactomePA package^38^ to obtain the enriched pathways for genes in the disease module with p-values lower than 0.05 cutoff adjusted by False Discovery Rate. Then, for each disease considered in this study, we extracted the disease-associated genes using the DisGeNET^39^ database and calculated the number of enriched pathways with at least two disease-associated genes. We found that the disease module calculated by RFIM shows the highest number of enriched pathways for five (six) diseases integrated with STRING (iRefīndex) in the disease module detection (see **Fig.5**) and this finding is valid if we counted the number of enriched pathways with at least four disease-associated genes (see **Fig.S2**).

**Figure 5:**
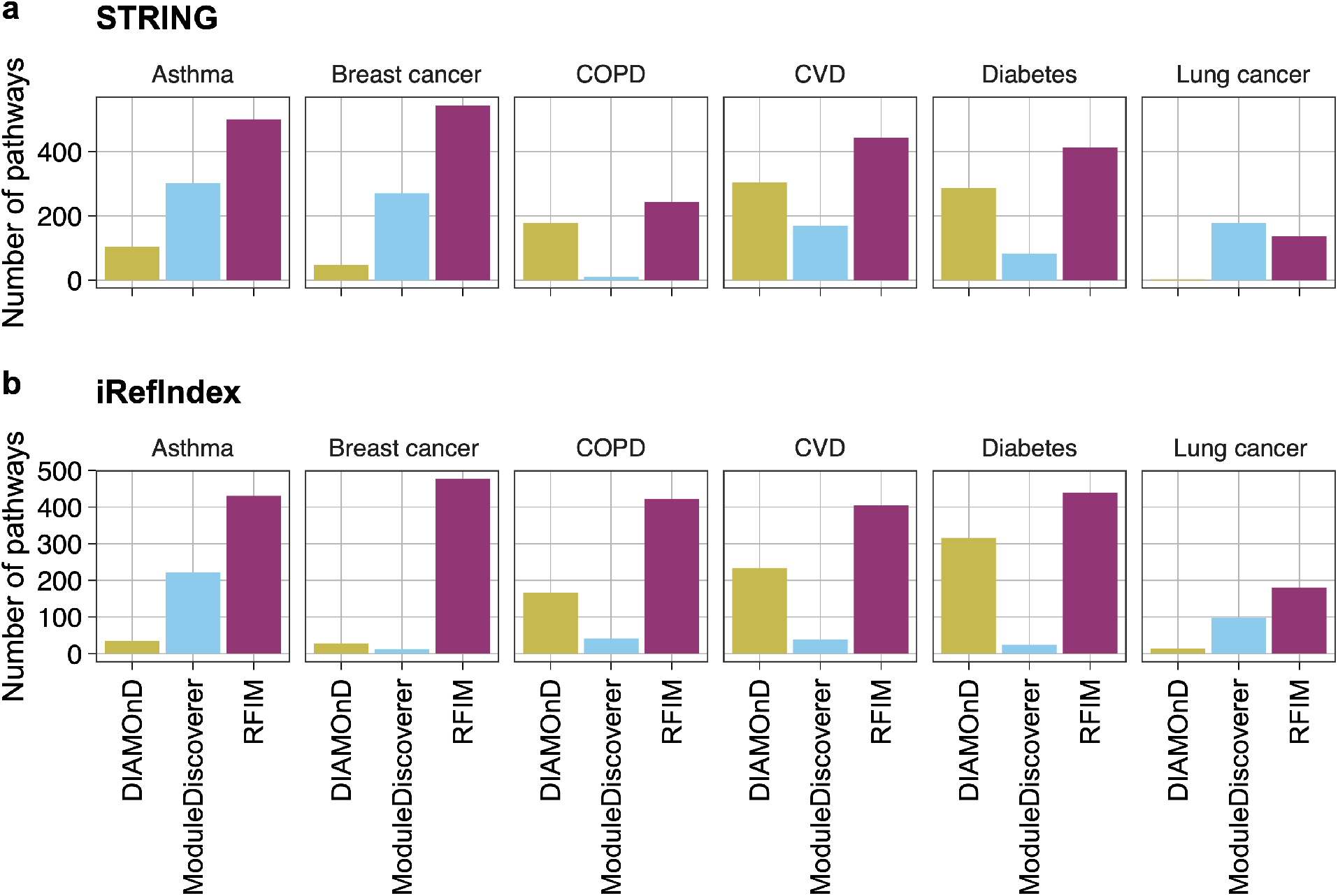
Enrichment analysis for genes in disease module detected from integrating the GWAS with two interactomes for each phenotype and method. We obtained the enriched pathways for disease genes using the ReactomePA package^38^ whose p-values are lower than 0.05 cutoff adjust by False Discovery Rate (FDR). Then, we extracted the disease-associated genes of each phenotype using the DisGeNET^39^ database and calculated the number of enriched pathways with at least two disease-associated genes.

### Robustness of disease module detection methods to the incompleteness of the interactome

The phenotype-specific disease module is apparently determined by both interactome and gene-wise p-values. Yet, the interactome is highly incomplete, covering a small fraction of known PPIs. To investigate the sensitivity of each disease module detection method to the incompleteness of interactome, we calculated the overlap between the disease modules for different levels of incompleteness of the underlying interactome. We firstly applied the disease module into the original interactome. Then, a fraction of PPIs randomly selected from the original interactome were removed. Finally, we performed the disease module detection algorithms again on the perturbed interactome and compared the fraction of overlapping genes in the union genes of the disease module identified from the original and perturbed interactome, respectively. Fig.6 shows that RFIM is more robust than other two methods, regardless of the fraction of PPIs have been removed.

**Figure 6:**
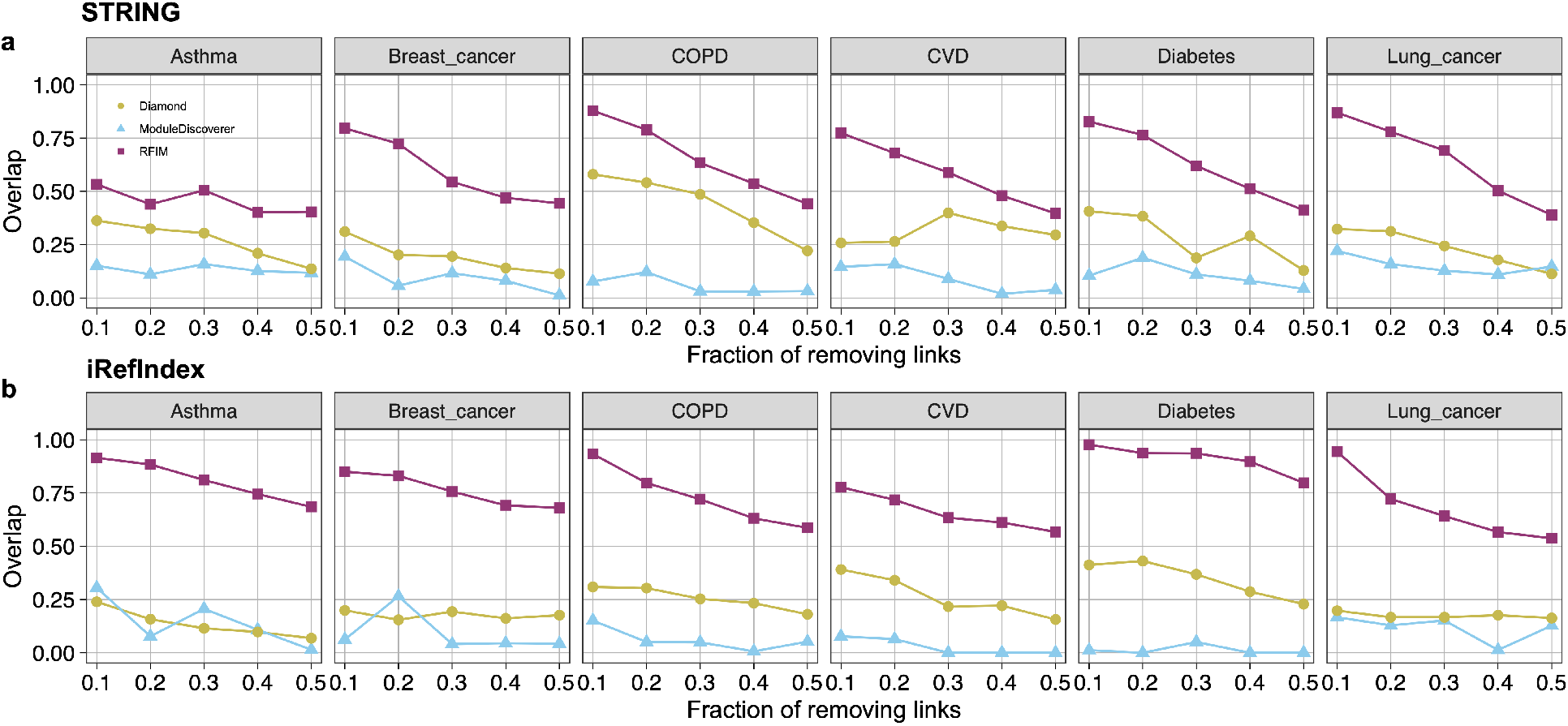
Sensitivity of disease module detection to the interactome. For each interactome, STRING (a) or iRefIndex (b), we firstly applied each disease module into the original interactome. Then, a fraction of PPIs randomly selected from the original interactome were removed. Finally, we performed the disease module detection algorithms again using perturbed interactome and compared the fraction of overlapping genes in the union genes of the disease module identified from the original and perturbed interactome, respectively.

### Disease module and enriched pathways

We visualized the structure of the disease module identified for asthma using our RFIM approach and the STRING interactome. We found that there are several sub-modules, each of which includes at least one gene that is significantly associated with disease (based on GWAS gene-wise p-value<0.001, see **Fig.7a** red nodes). The disease module apparently contains many genes that are not significantly associated with the disease (i.e., with GWAS gene-wise p-value>= 0.001, see **Fig.7a** blue nodes). Those genes are “absorbed” into the disease module due to their direct connection with those genes with very low GWAS p-values. We further performed the functional enrichment analysis of genes in the disease module using gprofiler2^40^. We found that these genes in the disease module are associated with many functional terms (see **Fig.7b**). Finally, we showed the top-10 enriched KEGG pathways (with lowest p-values in Fig.7b) enriched with genes in the disease module (see **Fig.7c**). We found that these pathways are potentially quite relevant to asthma. For example, proteasome inhibition is a novel mechanism to combat asthma^41^; Spliceosome are involved with the development of severe asthma^42^ and Notch signaling links embryonic lung development and asthmatic airway remodeling^43^. See **Fig.S3** and **S4** for the disease modules and functional terns of other disease phenotypes.

**Figure 7:**
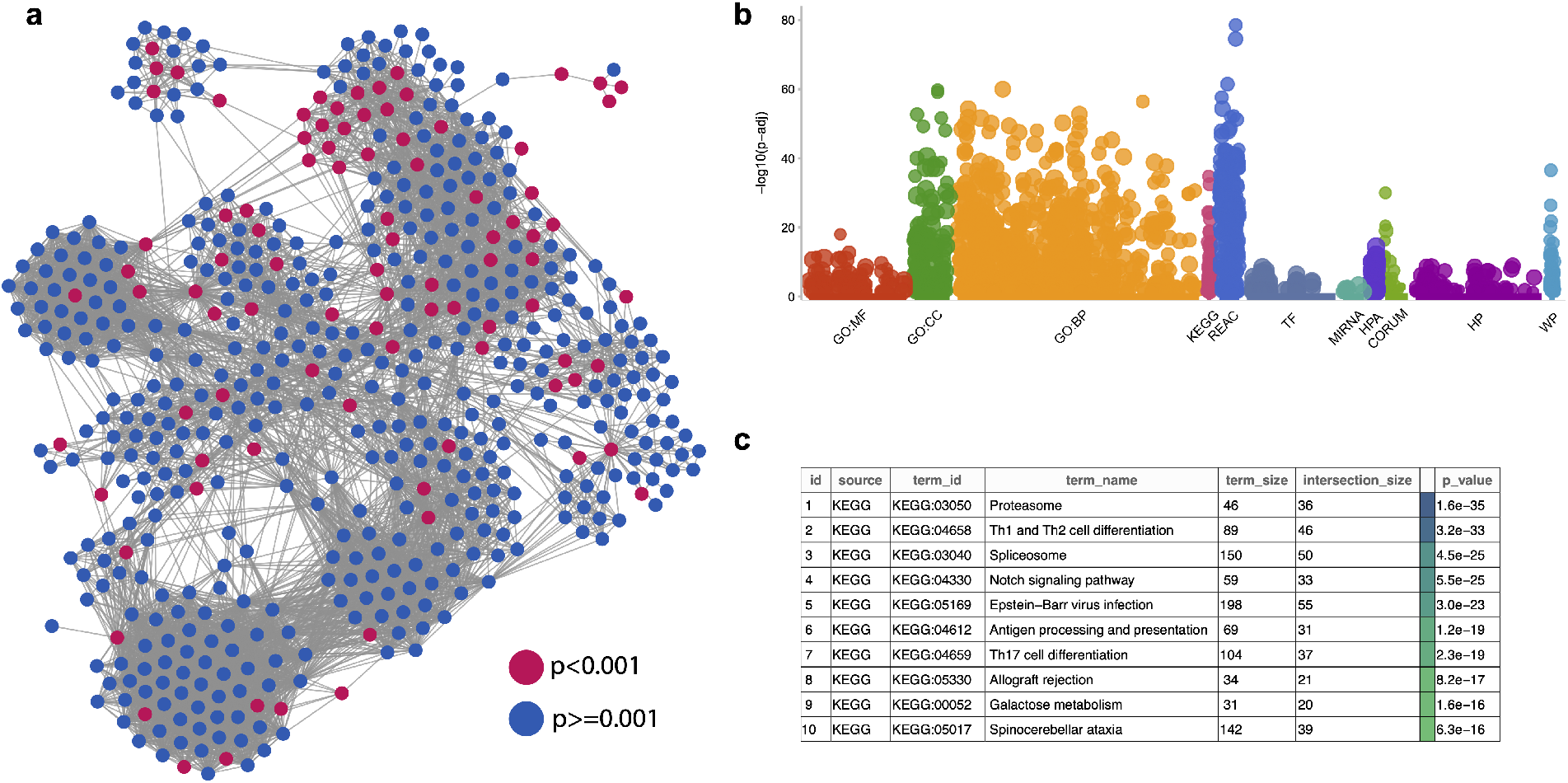
Sub-modules in the asthma disease module and the enriched functional terms. **(a):** Genes in the asthma disease module form several sub-modules. Genes with p-values lower (or higher) than 0.001 are colored with red (or blue), respectively. **(b):** Enriched functional terms of genes in disease module of asthma using gprofier2^40^. **(c):** Top-10 enriched KEGG pathways of genes in the asthma disease module using STRING.

### Literature validation for SNPs/genes in disease modules identified from GWAS

Finally, we explored the biological relevance of these genes belonging to the disease module. To narrow down the validation space, we focused on the common genes simultaneously identified by all three methods: RFIM, DIAMOnD and ModuleDiscoverer. We anticipate that those common genes are more important than genes identified by a single method. Interestingly, we found that those common genes were typically not significantly associated with the disease (based on the GWAS gene-wise p-values >= 0.001 (see **Table 2**). Yet, almost all these genes have been shown to be relevant for the particular disease. For example, for COPD, pulmonary proteasome expression and activity are downregulated and inversely correlated with lung function^44^. For asthma, the gene *IL9* has been implicated as an essential factor in determining susceptibility to atopic asthma and a therapeutic target for asthma^45^. For lung cancer, *CD74* has been associated with many tumor cells, i.e., non-small and small cell lung cancer^46,47^. And as a component of the HLA class-I complex, *B2M* has been shown to have recurrent inactivation in lung cancer. This was proposed to be an acquired mechanism for avoiding tumor immune recognition.^48^ For CVD, *APOC3*, an important regulator of TG homeostasis, is beneficially associated with decreased CVD risk^49–51^. These results imply that the genes in the disease module can be potentially associated with specific diseases despite their non-significance based on the current GWAS.

**Table 1.** Summary of GWAS studies used in the disease module detection.

| Phenotype | # Genes | # Cases | # Controls | Ref |
| --- | --- | --- | --- | --- |
| COPD | 17,866 | 15,256 | 47,936 | 24 |
| CVD | 17,866 | 60,801 | 123,504 | 25 |
| Diabetes | 17,894 | 11,645 | 32,769 | 26 |
| Asthma | 17,260 | 8,216 | 201,592 | 23 |
| Breast cancer | 17,215 | 5,552 | 89,731 | 23 |
| Lung cancer | 17,261 | 4,050 | 208,403 | 23 |

**Table 2.**
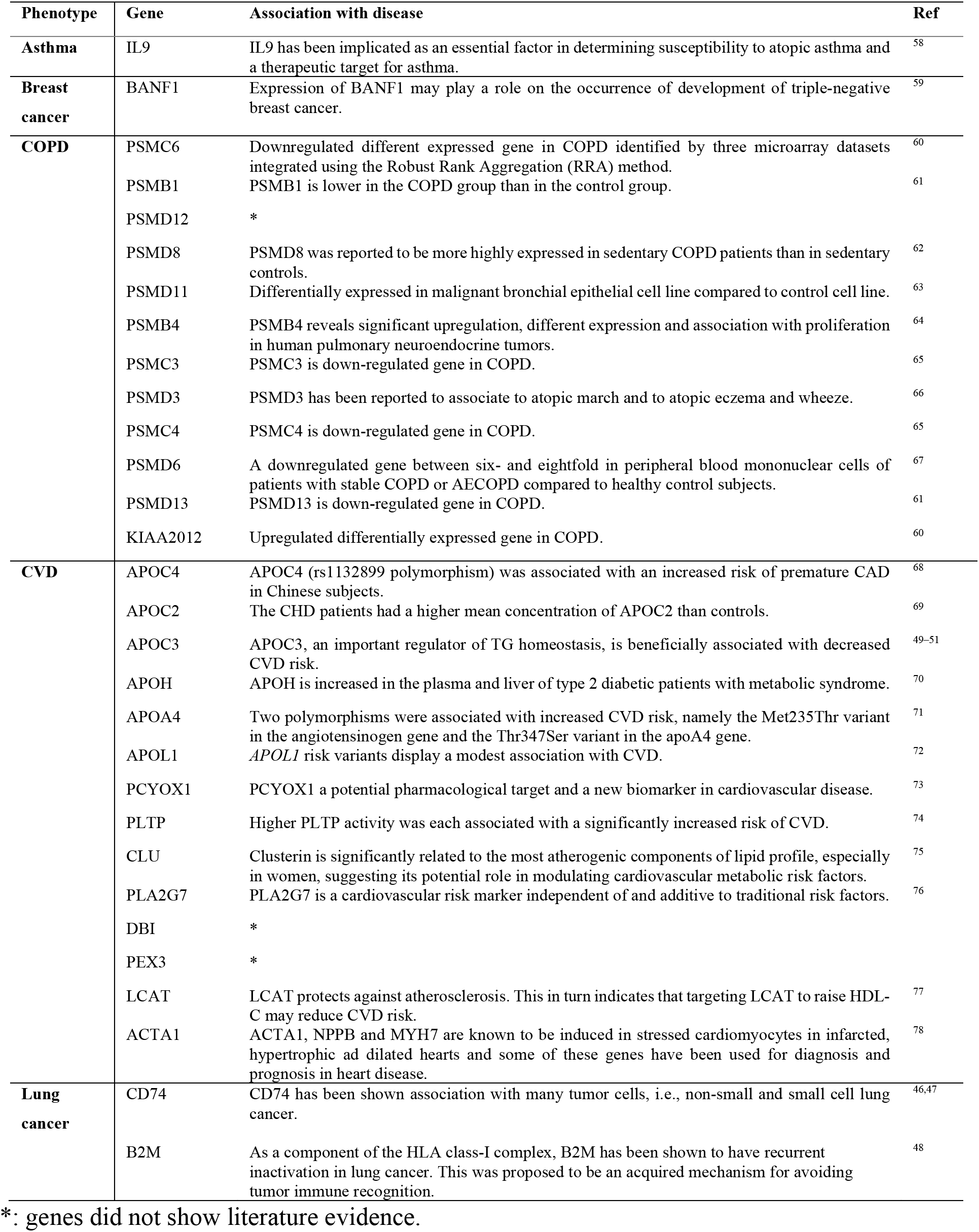
Biological relevance of genes simultaneously identified by three disease module detection methods.

## DISCUSSION

A complex disease can be linked to a well-defined neighborhood of the cellular molecular interactome, referred to as the disease module. Detecting the disease module allows us to identify driver genes that are heavily involved in the disease. Those driver genes do not necessarily have significant p-values in GWAS or other omics data, and hence cannot be easily identified without leveraging the human interactome. Here, we developed a novel disease module detection method based on a statistical physics model RFIM, which is both mathematically rigorous and computationally efficient. We applied our method to analyze six GWAS datasets, covering well-defined complex disease phenotypes. We found that the disease modules identified by RFIM display higher connectivity and more enriched pathways than the other existing methods. Furthermore, we found literature evidence that those genes in the disease module (especially those common genes identified by different methods) have significant associations with the specific disease.

Our RFIM approach can be naturally used to integrate other omics data, e.g., gene expression and DNA methylation, with the interactome. We admit that applying a disease module detection method via integrating omics data and the human interactome sometimes requires an annotation, which might lead to inaccuracy. For example, GWAS signals^32,52–54^ do not provide precise localization to a gene and the affected gene could be located at great genomic distance. Yet, we expect that using nearby genes is accurate enough to create meaningful disease modules.

## METHODs

### Calculation of the *M*(*H*) curve of RFIM

To calculate the *M*(*H*) curve of RFIM, we first need to calculate the exact ground state of the system at an arbitrary applied external field *H*. This is the basic step of calculating the *M*(*H*) curve, i.e., the ground state evolution for varying *H*. Fortunately, there is a well-known mapping of the ground state problem of RFIM to a min-cut/max-flow problem in combinatorial optimization. The mapping and the so-called push-relabel algorithm for the min-cut/max-flow problem has been well described in the literature^22^. The run time of the push-relabel algorithm^22^ scales as *O*(*N*^4/3^) with *N* the system size (number of nodes). The *M*(*H*) curve can be calculated with the method reported in [^49,50^]. It is essentially based on the fact that the ground state energy is convex up in *H*, which allows for estimates of the fields *H* where the magnetization jumps (called “avalanches”) occur. This algorithm finds steps by narrowing down ranges where the magnetization jumps with an efficient linear interpolation scheme^55^. Applying the no-passing rule (i.e., flipped spins can never flip back with increasing *H*) can further speed up the calculation of the RFIM *M*(*H*) curve^31^.

### Datasets

#### Interactome

For the human interactome, we used the STRING^33^ (9606.protein.links.v11.5.txt) database (http://string-db.org), which aims to provide a critical assessment and integration of PPIs, including both direct (physical) and indirect (functional) associations. We removed the PPIs whose normalized scores are lower than 0.7. The STRING Ensemble protein id (ENSP) is transformed into preferred gene name (e.g., entrez gene ID) using the protein information (9606.protein.info.v11.5.txt) downloaded from the STRING database. To examine the impact of different interactomes on the disease module detection methods, we also employed another interactome iRefIndex^34,35^ from the dataset extracted using R package cisPath^57^ (version 1.28.0).

#### GWAS

We used MAGMA^29^ (Multi-marker Analysis of GenoMic Annotation) to analyze the GWAS data. In MAGMA, SNPs were annotated to the genes based on dbSNP version 135 SNP location and NCBI 37.3. Two gene annotations were used in MAGMA. Only SNPs located between a gene’s transcription start and stop sites were annotated to that gene for the main analyses and a 10 kilobase window around each gene was made in additional annotation^29^. The GWAS data of CVD was downloaded from http://www.cardiogramplusc4d.org/data-downloads/ (version 2015). The GWAS data of diabetes was downloaded from http://diagram-consortium.org/downloads.html (version 2016). The GWAS data of COPD was collected in Ref [24], which combines 26 cohorts containing 63,192 individuals (15,256 COPD cases and 47,936 controls). The p-values and annotations of SNPs for CVD, diabetes and COPD were obtained using MAGMA. For the asthma, breast cancer and lung cancer data sets, the p-values of SNPs were directly downloaded from http://jenger.riken.jp/en/result and SNPs have been annotated to the genes in the summary results. The p-value of each gene in all datasets was obtained using mean *χ*^2^ statistic of SNPs included in the gene and a known approximation of the sampling distribution of the mean *χ*^2^ statistic^27,28^ using MAGMA.

### Existing disease module detection method used for comparison

There have many existing phenotype-specific disease module detection methods. For example, the DIseAse Module Detection (DIAMOnD) algorithm^18^ starts from a set of seed genes and prioritizes the other proteins of the interactome for their putative disease relevance. Therefore, the identified disease module might significantly rely on the initial seed genes. Another method, DysrEgulated Gene set Analysis via Subnetworks (DEGAS) detects the subnetworks in which multiple genes are dysregulated in the cases^15^. In the first step, the set of genes that are dysregulated when compared to controls are identified. Then, DEGAS identifies the smallest subgraph that contains at least *k* genes from each of those sets, except for up to *l* outlier sets from which fewer genes can be present. Parameters *k* and *l* are used to control the number of genes affected in the pathways in each individual and the number of allowed outliers, respectively, and several heuristics and algorithms were proposed to find these minimally connected subnetworks. In this work, we compared our proposed RFIM method with existing disease module detection methods incorporated in R package MODifieR^14^ which can simultaneously use the PPI network and gene-wise p-values. (1) ModuleDiscoverer^17^. ModuleDiscoverer uses a randomization heuristic-based approximation of the community structure based on maximal clique enumeration problem. (2) DIAMOnD^18^. DIAMOnD starts from a set of seed genes and prioritizes the other proteins of the interactome for their putative disease relevance. These methods are originally designed for gene expression data. The number of genes in the disease module in DIAMOnD set to be 500. We used the gene-wise p-value from GWAS as the input to ModuleDiscoverer and DIAMOnD.

---

**Figure S1:**
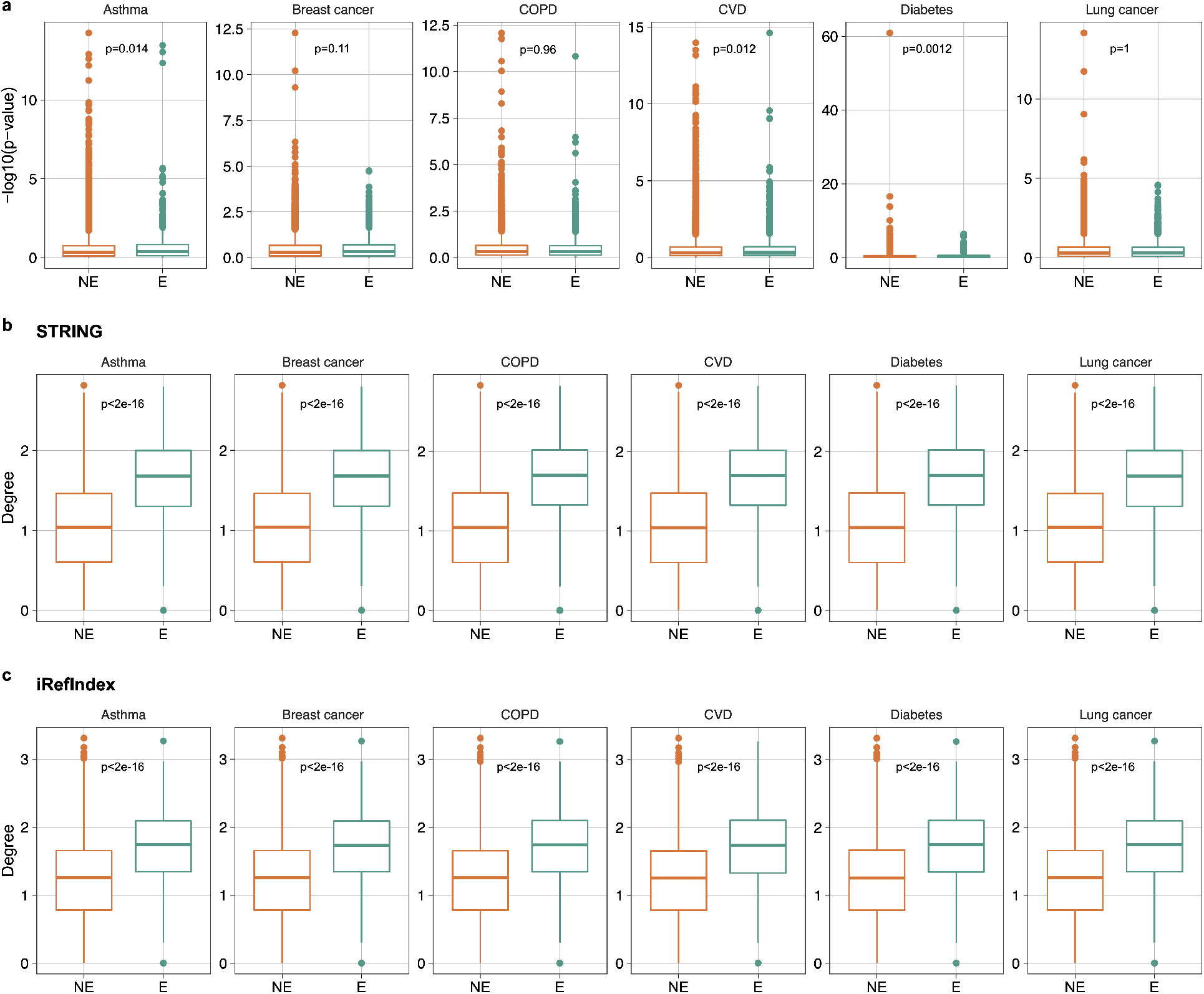
Correlation between gene essentiality, gene-wise p-value and degree. **(a):** Gene-wise p-value (in −log10 scale) distribution between essential and non-essential genes. **(b):** Gene degree (in log10 scale) distribution between essential and non-essential genes of STRING interactome. **(c):** Gene degree (in log10 scale) distribution between essential and non-essential genes of iRefIndex. Essential genes are defined as genes whose CS < −0.1 and corrected p< 0.05 in the KBM7 cell line. Significance level between essential and non-essential group were calculated using Wilcox test.

**Figure S2:**
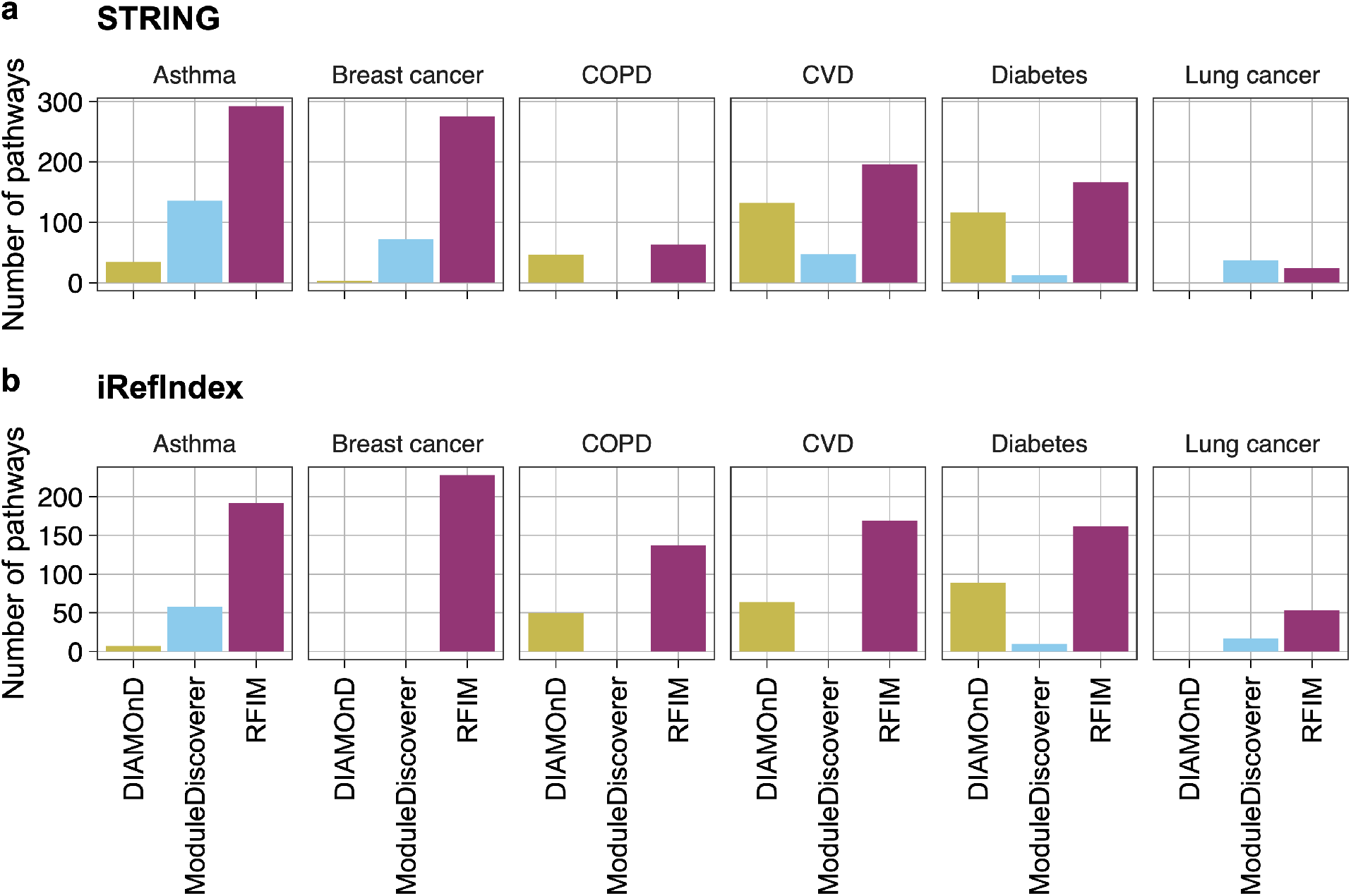
Enrichment analysis for genes in disease module detected from integrating the GWAS with four interactomes for each phenotype and method. We obtained the enriched pathways for disease genes using the ReactomePA package^38^ whose p-values are lower than 0.05 cutoff adjust by False Discovery Rate (FDR). Then, we extracted the disease-associated genes of each phenotype using the DisGeNET^39^ database and calculated the number of enriched pathways with at least two disease-associated genes.

**Figure S3:**
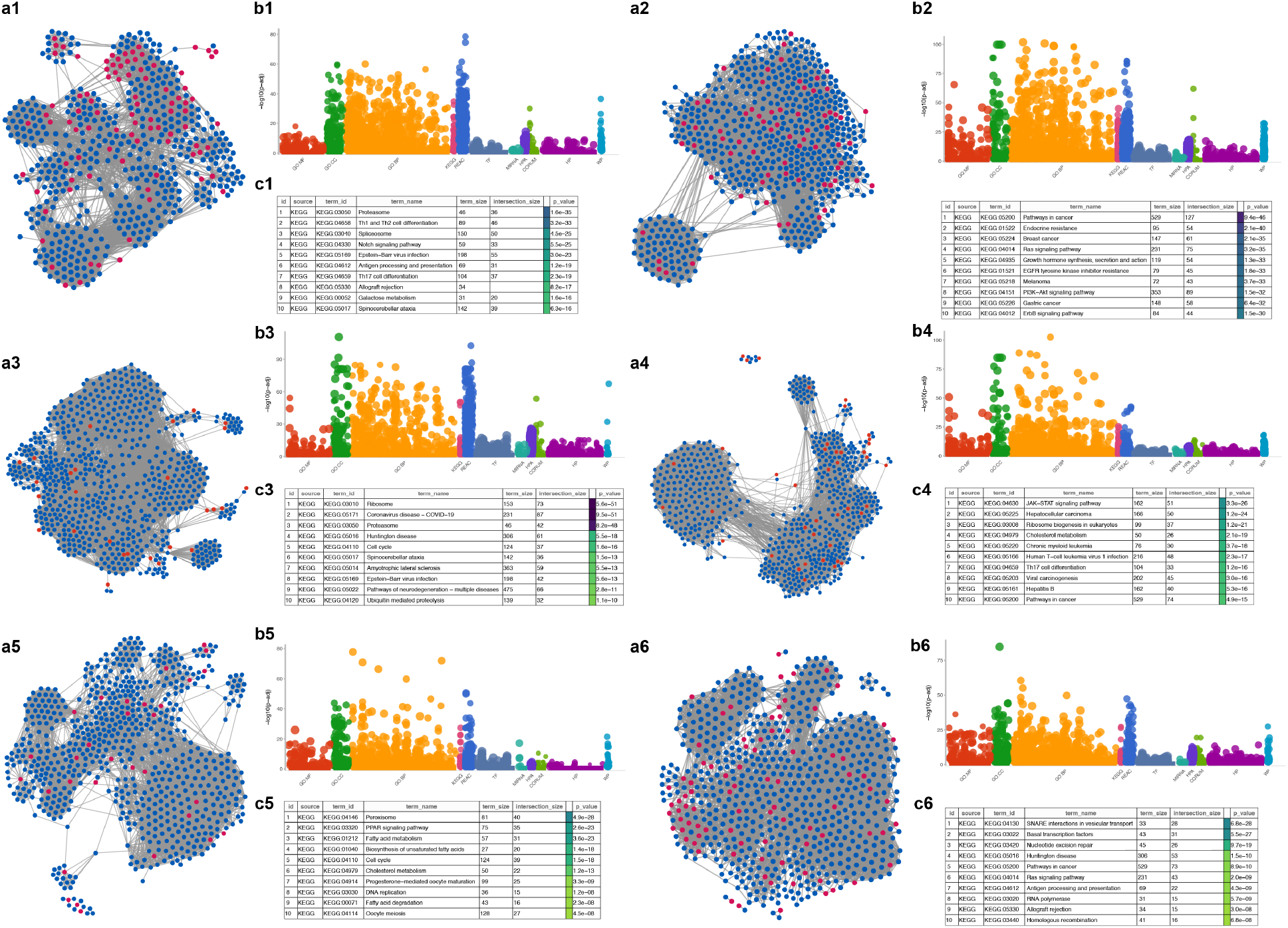
Subnetworks of genes in the disease module and enriched functional terms using STRING interactome. **(a1-a6):** Subnetwork of genes in disease modules of asthma, breast cancer, COPD, CVD, diabetes and lung cancer. Genes with p-values lower (higher) than 0.001 are colored with red (blue). **(b1-b6):** Enriched functional terms of genes in disease modules of asthma, breast cancer, COPD, CVD, diabetes and lung cancer using gprofier2^40^. **(c1-c6):** Top-10 enriched KEGG pathways of genes in the disease modules of of asthma, breast cancer, COPD, CVD, diabetes and lung cancer.

**Figure S4:**
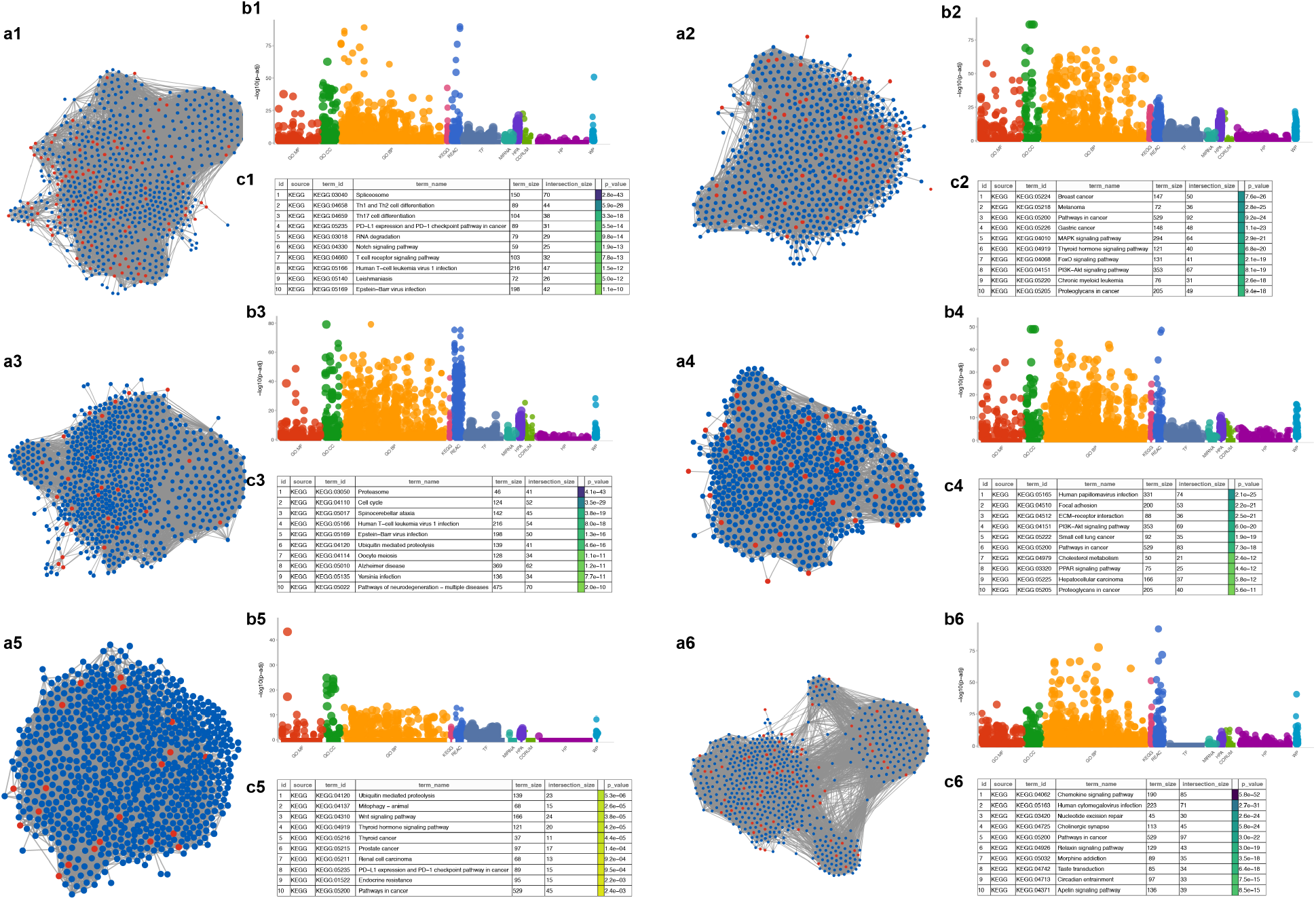
Subnetworks of genes in the disease module and enriched functional terms using iRefIndex interactome. **(a1-a6):** Subnetwork of genes in disease modules of asthma, breast cancer, COPD, CVD, diabetes and lung cancer. Genes with p-values lower (higher) than 0.001 are colored with red (blue). **(b1-b6):** Enriched functional terms of genes in disease modules of asthma, breast cancer, COPD, CVD, diabetes and lung cancer using gprofier2^40^. **(c1-c6):** Top-10 enriched KEGG pathways of genes in the disease modules of asthma, breast cancer, COPD, CVD, diabetes and lung cancer.

